# NOD2 activation reprograms infiltrating inflammatory monocytes in the Zika virus infected CNS to maintain neural correlates of learning and memory

**DOI:** 10.64898/2026.05.19.726300

**Authors:** Jeremy D. Hill, Alice H.W. Dong, Joshua Liu, Marina Diniz Barbezani, Tallulah S. Andrews, Robyn S. Klein

## Abstract

Zika virus (ZIKV) encephalitis induces cytokine-mediated cognitive deficits which persist long-term. Here, we determined if NOD2-mediated conversion of monocytes from Ly6C^Hi^ inflammatory to Ly6C^Lo^ anti-inflammatory phenotypes during ZIKV encephalitis preserves neural correlates of learning and memory within the hippocampus. Short-term administration of the NOD2 agonist, muramyl dipeptide (MDP), during peak ZIKV infection prevents synapse elimination and loss of adult hippocampal neurogenesis, without impacting CNS virologic control. Transcriptomic analyses of forebrain immune cells in MDP-treated mice revealed functional modulation of infiltrated monocytes and T cells, reducing their expression of pro-inflammatory cytokines, with limited effects on microglia, compared to controls. Notably, NOD2 activation in peripheral immune cells alone balances innate immune signals, preserving synapses, and increasing macrophage phagocytic capacities that do not target synapses. Our findings identify infiltrating Ly6C^Hi^ monocytes as key drivers of long-term cognitive dysfunction following ZIKV encephalitis and as potential therapeutic targets for limiting synapse loss.

**Highlights:**

- NOD2 activation via MDP phenotypically shifts monocyte subsets from inflammatory to anti-inflammatory during the acute phase of ZIKV encephalitis.
- Short-term MDP administration increases phagocytic machinery and reduces inflammatory cytokine/chemokine production within macrophage populations.
- Synapse elimination is attenuated within the hippocampus of ZIKV-infected animals treated with MDP.
- MDP derived effects are mediated through peripherally derived immune cells.

## Introduction

Macrophages play a crucial dual role in viral encephalitis. They act as essential defenders by neutralizing viruses and modulating immune responses of infiltrating T cells. Conversely, their overactivation can induce severe neuroinflammation, driving persistent activation of microglia, astrocytes, and cytotoxic CD8 T cells^1,2^. Nucleotide-binding oligomerization domain 2 (NOD2) is a cytosolic pattern recognition receptor belonging to the NOD-like receptor (NLR) family, which plays a central role in innate immune sensing of bacterial peptidoglycan-derived muramyl dipeptide (MDP)^3–5^. Highly expressed in monocyte and macrophage populations, activation of NOD2 by MDP orchestrates multiple downstream responses through activation of the NFκB and MAPK signaling pathways, promoting regulation of autophagy responses and enhanced antigen presentation^6–11^. Emerging evidence indicates that NOD2 interacts with multiple innate immune pathways and acts as a regulator of macrophage polarization. NOD2 stimulation via MDP can induce distinct pro-inflammatory (interleukin 1 beta [IL-1β], interleukin 6 [IL-6], tumor necrosis factor alpha [TNFα]) or anti-inflammatory cytokine (interleukin 10 [IL-10]) profiles demonstrating that the same NOD2 stimulus can generate qualitatively different outputs^12^. Leveraging this dual capacity to modulate immune responses, NOD2 activation may be harnessed to reprogram maladaptive immune responses and restore homeostasis in the context of chronic inflammatory diseases.

The therapeutic administration of MDP represents a shift away from traditional immunosuppression toward the induction of immunological tolerance. In a clinical study of rheumatoid arthritis, MDP administration induced significant increases in CD4^+^/CD25^+^/FoxP3^+^ Tregs and transforming growth factor beta (TGFβ) in blood samples which were associated with reductions in joint swelling and stiffness, suggesting a shift to an anti-inflammatory environment^13^. In animal studies, chronic, low-dose MDP treatment phenotypically shifts macrophages away from an inflammatory M1 phenotype to an M2b signature with high expression of MHCII and IL-10^7,14^. This “regulatory” phenotype attenuates the recruitment of inflammatory cells and initiates repair mechanisms to limit cellular damage^15,16^. This altered phenotype also influences subsequent pattern recognition receptor activation; in response to challenge with a high dose of LPS, MDP-treated macrophages produce significantly less pro-inflammatory mediators and oxidative byproducts and more IL-10 than non-MDP treated cells^14^. While these findings highlight the potential for MDP-NOD2 based therapeutic strategies to mitigate chronic inflammation, it is unknown whether short-term treatment in the context of infectious diseases is safe and impacts long-term sequelae, including cognitive impairment post viral encephalitis.

Recovery from acute encephalitis induced by mosquito-borne, neurotropic orthoflaviviruses, such as Zika virus (ZIKV), is associated with significant long-term cognitive sequelae which may persist and worsen for years despite viral clearance^17–19^. The formation and consolidation of new memories occurs primarily within the hippocampus, and relies on the integrity of trisynaptic circuits between the entorhinal cortex, hippocampal dentate gyrus (DG) and Cornu Ammoni (CA) regions^20,21^. Spatial learning also depends on rates of adult neurogenesis, which occurs via generation of new neurons from neural stem cells within the subgranular zone of the DG^22,23^. New neurons form synaptic connections with existing neuronal networks, supporting normal memory formation^23–26^. During the acute phase of viral encephalitis, infiltrating T cells and activated macrophages/microglia produce antiviral/inflammatory cytokines which act on infected neurons and glial cells to induce antiviral gene expression, inhibit viral replication, and enhance antigen presentation^27–31^. Despite effective viral clearance by virus-specific CD8^+^ T cells, their persistence as resident memory CD8 T cells (CD8 T_RM_) chronically activates resident microglia and astrocytes, leading to prolonged synapse elimination and dysregulation of adult neurogenesis within the hippocampus^32–37^. Thus, exposure to neurotropic pathogens is now included as a risk factor for the development of neurodegenerative diseases, further indicating a significant need to better understand inflammatory mechanisms triggered during acute orthoflavivirus infections and their contribution to persistent neurologic sequelae^38^.

In this study, we demonstrate that NOD2-mediated conversion of inflammatory Ly6C^Hi^ monocytes to an anti-inflammatory Ly6C^Lo^ phenotype during ZIKV encephalitis reduces neuroinflammation, increases phagocytic capacity of infiltrating monocyte/macrophage populations, prevents synapse elimination and limits loss of adult neurogenesis within the hippocampus. These effects are monocyte dependent as microglial and astrocyte populations showed limited alterations in function during MDP administration, as revealed by single cell RNAseq analyses. Moreover, bone-marrow chimeric studies utilizing NOD2-deficient mice show that macrophage phagocytosis and synapse preservation requires MDP-dependent NOD2 signaling only in peripheral myeloid-derived cells for therapeutic effects. These observations indicate that modulation of infiltrating Ly6C^Hi^ inflammatory monocyte phenotype via NOD2 activation protects against alterations in neural correlates of learning and memory within the hippocampus that underlie post-recovery cognitive sequelae.

## Results

### Administration of MDP converts blood and brain monocytes from Ly6C^Hi^ to Ly6C^Lo^ during acute ZIKV infection without impacting virologic control

To assess the specific influence of peripherally derived Ly6C^Hi^ inflammatory monocytes on neural correlates of learning and memory during ZIKV infection, we administered MDP (10mg/kg) or the inactive isoform of MDP (iMDP; 10mg/kg) i.p. to mice to modulate infiltrating monocytes from a Ly6C^Hi^ to Ly6C^Lo^ phenotype. Employing a previously published, recovery model of adult ZIKV encephalitis^34^ in which 8-10 week-old C57BL/6J (wild-type (WT)) mice are i.c. infected with ZIKV-Dakar (Dakar 41525, Senegal, 1984), we initiated MDP treatment at 7 days post-infection (dpi) (Fig.1A), which coincides with peak infection in our model^34^, a clinically relevant timepoint at which to begin treatment, and continued daily treatment until 10 dpi. Short-term administration of MDP did not alter survival rates, weight loss, encephalitic scores, or viral clearance compared with similarly infected, iMDP-treated controls (Fig. 1B-E). Flow cytometric analysis of blood at 10 dpi showed a significant phenotypic shift in the monocyte population of MDP-treated animals (∼20% Ly6C^Hi^ cells and ∼60% Ly6C^Lo^ cells) compared with iMDP-treated mice (60% Ly6C^Hi^ and ∼20% Ly6C^Lo^ cells) (Fig. 1F). The gating strategy used for flow analyses can be seen in Fig. S1. MDP had no effect on the percentages of CD4^+^ or CD8^+^ T cells within the blood at 10 dpi (Fig. 1G). Within the hippocampus (HPC), the same reciprocal shift in monocyte phenotype was observed at 10 dpi (Fig. 1H). MDP did reduce the total number of T cells in the HPC at this time-point, suggesting that MDP may impact their overall recruitment into the CNS (Fig. 1I). MDP also did not alter the number of P2RY12^+^ cells present in the hippocampus, indicating no effect on microgliosis (Fig. 1J). Notably, other methods for eliminating inflammatory monocyte infiltration during ZIKV infection, including antibody-based depletion targeting Ly6C (Fig. S2A) or Gr1 (Fig. S2B), or use of mice with global deletion of CCR2 exhibited off-target effects (Fig. S2C), reducing numbers and percentages of blood and brain CD8 T cells, associated with increased mortality with lack of virologic control. Overall, these data show that short-term treatment with MDP, beginning at peak infection (7 dpi) until 10 dpi, converts CNS infiltrating monocyte subsets from a Ly6C^Hi^ inflammatory to Ly6C^Lo^ anti-inflammatory phenotype without impacting viral clearance or survival during ZIKV encephalitis. Use of MDP also provides a robust tool to evaluate how CNS infiltrating, pro-inflammatory monocytes contribute to ZIKV-mediated alterations in neural correlates of learning and memory.

**Figure 1.**
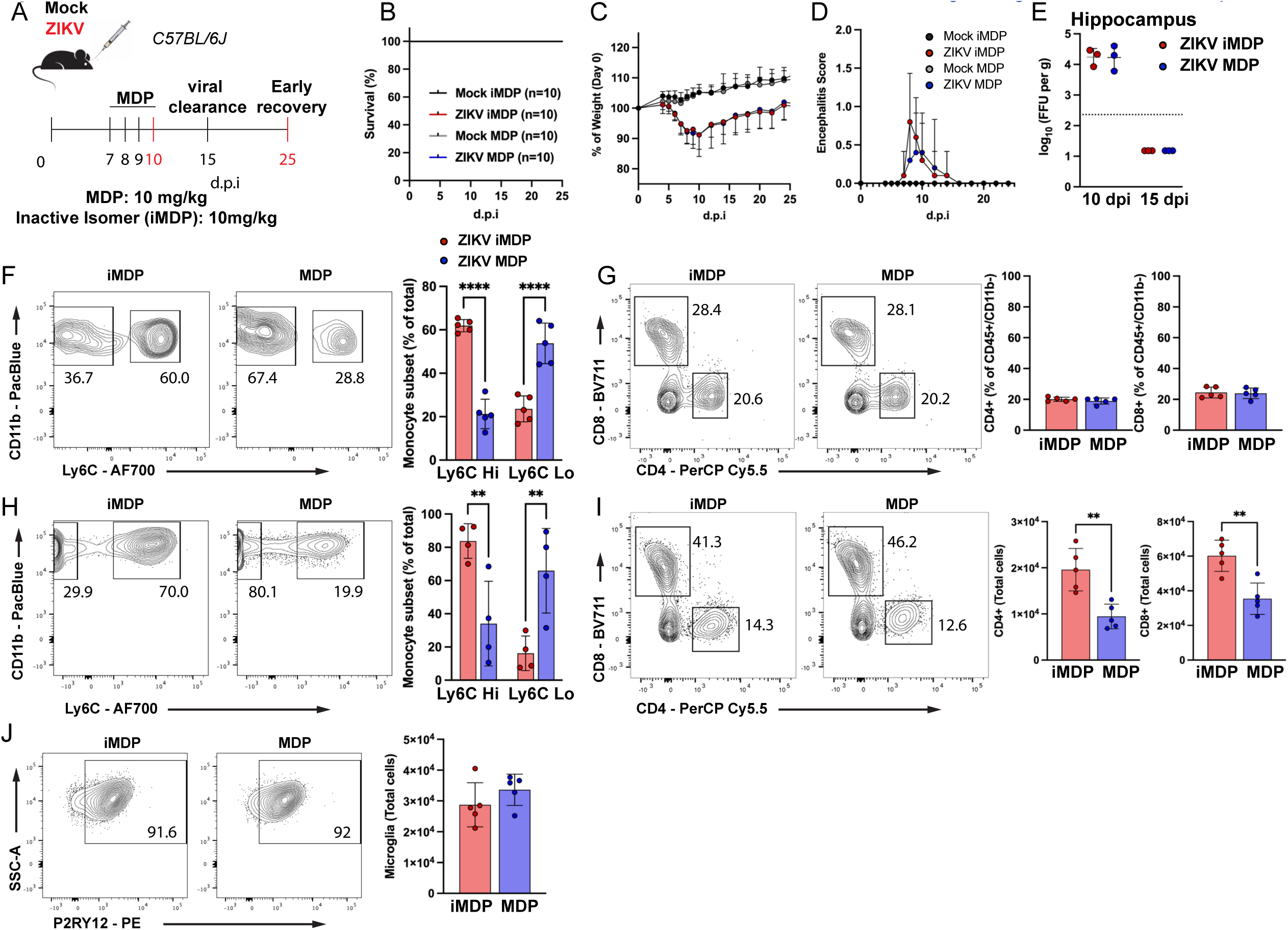
Administration of MDP converts blood and brain monocytes from Ly6C^Hi^ to Ly6C^Lo^ during acute ZIKV infection without impacting virologic control. 8 week-old C57BL/6J animals were inoculated intracranially with ZIKV-Dakar (1x10^4^ FFU) and underwent inactive isomer of MDP (10mg/kg) or MDP (10mg/kg) treatment. (A) Schematic representation of viral infection, MDP administration, and tissue collection timepoints. (B) Treatment with MDP did not induce lethality in 8-week-old ZIKV-infected mice. (C) ZIKV infection led to weight loss in infected animals. MDP treatment did not alter weight loss when compared to ZIKV-infected, inactive isomer-treated animals. Data is expressed as the percentage of Day 0 body weight. (D) ZIKV-infected animals showed increases in encephalitic score, which did not differ between inactive isomer and MDP-treated animals. Data were pooled from two independent experiments (n=10 animals per group) (E) MDP did not alter viral burden within the hippocampus at 10 or 15 days post infection. Viral burden was assessed via focus forming assay (n=3 per group).Dotted line on graph is limit of detection. (F-G) (F) Representative flow plots and analysis of Ly6C expression in CD11b+ monocytes, (G) CD4^+^ and CD8^+^ T cells within the blood at 10 dpi. MDP treatment (started at 7 dpi) lead to a switch from a Ly6C^Hi^ phenotype to Ly6C^Lo^. MDP treatment did not show alteration of the percent of CD4^+^ or CD8^+^ T cells as a proportion of CD45^+^/CD11b^-^cells within the blood at 10 dpi. Multiple experiments were conducted, representative results from one experiment are shown (n=5 per group in shown experiment). (H-J) Analysis of bone marrow derived monocytes, T cells, and microglia within the hippocampus at 10 dpi. (H) representative flow plots and analysis of Ly6C expression in CD45^Hi^/ CD11b^+^ monocytes within the hippocampus. CNS infiltrated monocytes continue to show a switch from a Ly6C^Hi^ phenotype to Ly6C^Lo^ due to MDP treatment. (I) flow cytometric analysis of CD4^+^ and CD8^+^ T cells within the hippocampus. MDP-treated animals showed significantly lower numbers of infiltrating CD4^+^ and CD8^+^ T cells when compared to iMDP-treated controls. (J) Representative flow plots and analysis of total numbers of microglial cells (CD45^Mid^/CD11b^+^/P2RY12^+^ cells) were unchanged due to MDP treatment within the hippocampus. Multiple experiments were conducted, representative results from one experiment are shown (H, n=4 per group, I,J, n=5 per group in shown experiment). All data are represented as mean ± s.d. and were analyzed by two-way (F, H) ANOVA or student’s t-test (G, I, J). \*\**p* < 0.01, \*\*\*\**p* < 0.0001.

### MDP treatment induces a macrophage phenotypic shift and upregulates macrophage phagocytic markers in the ZIKV-infected HPC

To define how NOD2 activation influences transcriptional signatures of mononuclear cells within the ZIKV-infected forebrain, we performed scRNA-seq analyses on forebrain tissues derived from iMDP- versus MDP-treated ZIKV-infected mice at 10 dpi. As done previously^39^, we utilized an isolation protocol to enrich for leukocyte populations. Forebrains were collected from 6 animals for each treatment, and cells from 3 brains per treatment were pooled to generate 2 biological replicates for each treatment. After quality control, filtering and data normalization (see Methods), a total of 20489 cells were used for downstream analyses. Clustering of selected major cell types revealed 12 distinct cell populations distributed across both treatment groups (Fig 2A). The cluster of monocyte-derived macrophages (MDMs) exhibited the highest numbers of differentially expressed genes (DEGs), while transcriptomic changes in microglia, T cell subtypes, and other immune cells were modest in comparison (Table S3). Significantly decreased DEGs in monocytes derived from the forebrain of MDP-treated mice compared with forebrain derived from iMDP-treated animals included genes involved in inflammatory cytokine/chemokine signaling (*Il1b, Ccl4, Ccl5, Cxcl9, Cxcl10*) (Fig 2B), which critically regulate leukocyte infiltration. Importantly, MDP treatment led to significant upregulation of pathways associated with anti-inflammatory M2 phenotype differentiation (*Arg1, Grn, Chil3*) and phagocytosis/lysosome-related genes (*Trem2, Mertk, Ctsd, Ctsl, Grn, Lamp1, Fcgr3)* (Fig 2B, C), indicating that MDP treatment induces infiltrating MDMs to adopt a more M2b-like phenotype with enhanced phagocytic machinery. Pathways associated with proinflammatory cytokine signaling, such as responses to IFNβ, IL-2 production, responses to lipopolysaccharide, and IL-17 signaling, were among the top 50 significantly downregulated pathways (Fig 2C), supporting a shift to a less inflammatory phenotype. Differential expression and pathway enrichment analyses on CD8 T cells similarly showed decreased chemokine signalling (*Ccl4*) and chemotaxis pathways (Fig. 2D, E). Which is predicted to result in decreased CCL4-CCR5 and CXCL10-CXCR3 interactions between CD8 T cells and macrophages (Fig. 2F). However, these CD8 T cells were shifted to a more effector-like phenotype with increased expression of genes associated with cell cycle progression (*Cdk6*), survival (*Bcl2*), and activation (*Gzmb, Klrd1, Tnfrsf9, Tnfrsf18, Lgals1*) (Fig. 2D).

**Figure 2.**
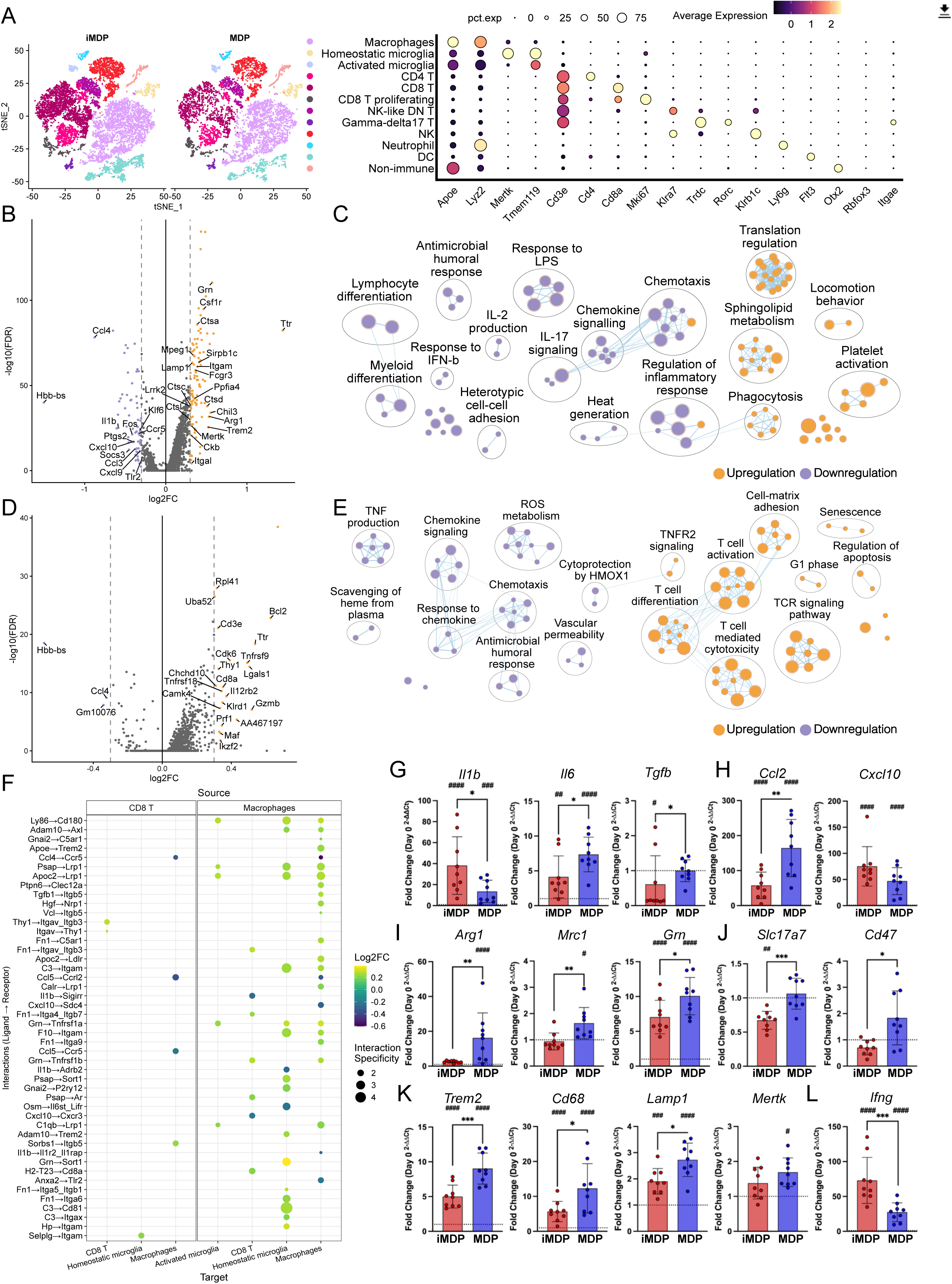
Pro-inflammatory immune responses are altered in monocyte-derived macrophages and T cells in the forebrain of ZIKV-infected mice after treatment with MDP. Mice were intracranially infected with ZIKV, then treated with i.p. injections of either inactive isomer (iMDP) or MDP (n = 2). At 10 dpi, single-cell RNA sequencing was then performed on all cells from the forebrain of these mice, with two forebrains pooled into a sample. (A) Left, tSNE representation of all cells in the forebrain of ZIKV-infected mice after treatment with iMDP or MDP, colored by cell type. Right, normalized expression of characteristic genes in each cell type. (B, D) Differentially expressed genes (DEGs) among cells in the (B) monocyte-derived macrophage and (D) CD8+ T cell clusters. Genes were defined as differentially expressed if the log2-transformed fold-change was beyond 0.3 in magnitude and the adjusted p-value was less than 0.05. (C, E) Differentially expressed pathways among cells in the (C) monocyte-derived macrophage and (E) CD8+ T cell clusters. Overrepresentation analysis was performed using the significant DEGs from each cluster, and only the top 50 significantly upregulated or downregulated pathways were depicted (adjusted p value < 0.05). Each colored dot represents a single pathway, and all significant pathways were clustered using the MCL algorithm. Lines connecting colored dots depict genes shared between pathways. (F) Putative differential cell-cell communication among CD8+ T cells and myeloid cells. Putative ligand-receptor interactions and their corresponding interaction specificities (-log10(specificity rank)) were determined using cells isolated with iMDP-treated mice. Log2FC refers to the average log2-transformed fold-changes of the ligand and receptor in the corresponding cell type when comparing the expression in iMDP-treated mice to MDP-treated mice. Only interactions with Log2FC magnitude greater than or equal to 0.2 and specificity rank less than or equal to 0.05 are shown. (G-L) qPCR analysis of (G) macrophage-associated cytokines, (H) chemokines, (I) M2-associated markers, (J) neuronal-associated markers, (K) sensors for damage and lysosomal activation, and (L) CD8^+^ T cell-derived cytokines. Each dot represents an individual mouse (G-L, Day 0 n=6, iMDP n=9, MDP n=9). Data are presented as fold change of Day 0 samples (mean ± s.d.). Dotted lines on graphs represent fold change of Day 0 samples (1). Asterisks denote statistical significance between iMDP and MDP treatments (\**p*<0.05, \*\**p*<0.01, \*\*\**p*<0.001). Hash signs denote statistical significance between Day 0 and treatment (#*p*<0.05, ##*p*<0.01, ###*p*<0.001, ####*p*<0.0001). Statistical analysis was performed using one-way ANOVA with multiple comparisons (G-L).

Validation of macrophage cytokine mRNAs in hippocampi derived from separate cohorts of MDP- versus iMDP-treated, infected animals at 10 dpi confirmed reduced levels of *Il1b*, and increased *Il6*, *Tgfb*, and *Ccl2* in comparison to iMDP-treated animals (Fig 2G, H). MDP treatment also led to increased expression of markers of M2 macrophage phenotype conversion (*Arg1*, *Mrc1, Mafb, Grn*) and sensors for damage and lysosomal activation (*Trem2, Cd68,* and *Lamp1*) (Fig. 2I, K, Fig. S3A) with no overall changes in genes associated with the classical complement pathway (Fig. S3B). MDP also had no effect on genes associated with phagocytosis (*Axl*, *Maf*, *Sirpa*, Fig. S3A), cytokines and aspects of inflammasome signaling (*Il4*, *Il10*, *Nlrp3*; Fig. S3C), synaptic maintenance (*Dlg2*, *Dlg4*, *Grin1*, Fig. S3D), and activity-dependent transcription (*Fos,* Fig. S3E). Notably, MDP treatment attenuated loss of *Cd47* and *Slc17a7*, the latter of which is a marker of mature excitatory neurons which acts with Cd47 to protect synapses (Fig 2J). Analysis of antiviral CD8^+^ T cell-derived *Ifng* showed significantly lower expression in MDP-treated hippocampi as compared to iMDP (Fig 2L). These data suggest that MDP administration alters the phenotype of infiltrating monocytes into a more anti-inflammatory state, with increased phagocytic/lysosomal machinery, and attenuates ZIKV-induced reductions of signals that maintain synaptic proteins and plasticity.

### MDP-treated mice exhibit increased phagocytic myeloid cells during acute ZIKV infection

Prior studies have shown that the transition from acute neuroinflammation to the resolution phase is characterized by the upregulation of M2 macrophage phagocytosis markers (*Cd68*, *Lamp1*) and driven by the coordination of IL-4, IL-10, and TGFβ, which promotes the clearance of cellular debris and suppression of microglial activation^40–44^. Given similar findings in our transcriptomic data, we examined proteins associated with phagocytic myeloid cells via immunohistochemical (IHC) analysis at 10 dpi. Colocalization of CD68+ area within IBA-1+/Tmem119- cells in MDP-treated, ZIKV-infected animals was significantly increased compared to iMDP-treated, infected mice (Fig. 3A, B). This was accompanied by a significant increase in the number of macrophages found in MDP-treated ZIKV-infected animals as assessed by manual counting of IBA-1+/Tmem119- cells within the CA3 region of the HPC. Similar evaluation of animals during recovery (25 dpi) (Fig. 3C, D) revealed that only iMDP-treated, ZIKV-infected animals continued to exhibit elevated CD68+ area in Iba1+/TMEM119-negative cells compared to mock-infected controls, and the significant increase in macrophage numbers due to MDP had resovled. To assess whether increased CD68 was associated with enhanced lysosomal activity at 10 dpi, we immunostained for Lamp1 and observed similar increases within Iba1+ cells in MDP-treated, ZIKV-infected animals when compared to iMDP treatment (Fig. 3E, F). Overall, these data indicate that MDP treatment significantly increases phagolysosomal activity within Tmem119^-^ myeloid cells during ZIKV encephalitis.

**Figure 3.**
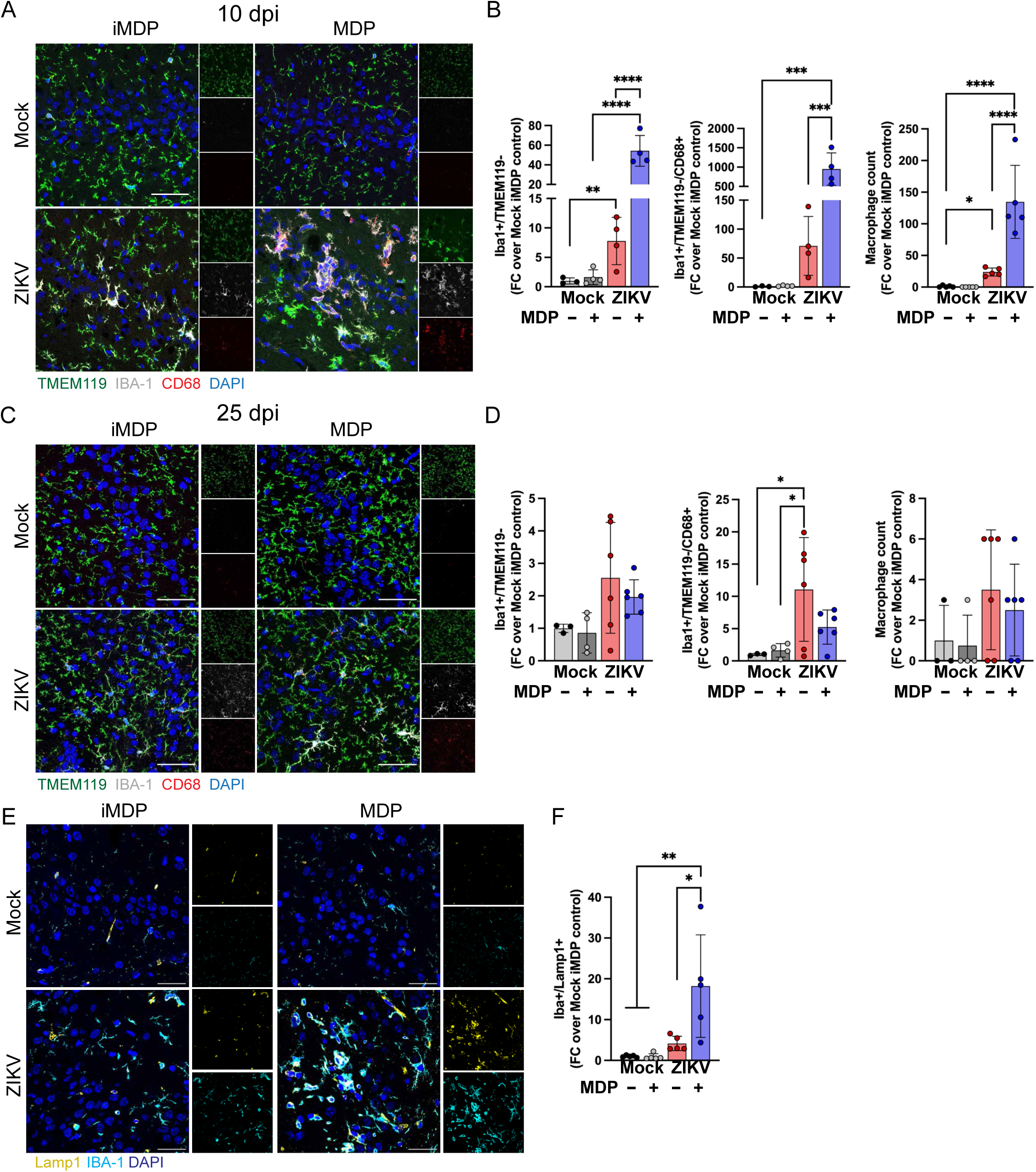
Administration of MDP increases lysosomal marker expression within infiltrating myeloid cells during the acute phase of ZIKV-induced encephalitis. (A, B), ZIKV infection induces increased expression of IBA-1 and CD68 in the hippocampus at 10 dpi, which is significantly increased by treatment with MDP. (A) Representative immunostaining of infiltrating myeloid cells (IBA-1+ [grey] /TMEM119-[green]) colocalized with CD68 (red) at 10 dpi with or without MDP treatment in mock or ZIKV-infected animals. Mock iMDP n=3, Mock MDP n=4, ZIKV iMDP n=4, ZIKV MDP n=4. (B) Quantification of IBA1+/TMEM119- area, IBA1+/TMEM119-/CD68+ area, and macrophage numbers at 10 dpi (displayed as fold change of Mock iMDP controls). (C) Representative immunostaining of infiltrating myeloid cells (IBA-1+ [grey] /TMEM119-[green]) colocalized with CD68 (red) at 25 dpi with or without MDP treatment in mock or ZIKV-infected animals. Mock iMDP n=3, Mock MDP n=4, ZIKV iMDP n=6, ZIKV MDP n=6. (D) Quantification of IBA1+/TMEM119- area, IBA1+/TMEM119-/CD68+ area, and macrophage numbers at 10 dpi (displayed as fold change of Mock iMDP controls). (E) Representative immunostaining of IBA1 (cyan) colocalized with Lamp1 (yellow) at 10 dpi. (F) Quantification of IBA1+/Lamp1+ area and displayed as fold change over Mock iMDP control (n=5 per group). Data were pooled from two independent experiments. Scale bars, 25 µm. Data are displayed as mean ± s.d. and were analyzed by two-way ANOVA, corrected for multiple comparisons. All panels, \**p*<0.05, \*\**p*<0.01, *\*\*\*p<* 0.001, \*\*\*\**p*< 0.0001.

### MDP treatment maintains post-synaptic termini during ZIKV encephalitis

In previous studies, we and others found that ZIKV encephalitis is associated with significant elimination of Homer1+ post-synaptic termini within the CA3 region of the HPC beginning at 7 dpi, followed by neuronophagia by 25 dpi^34,45,46^. Given that administration of MDP significantly reduced *Il1b* expression, increased phagocytic pathways, but had no effect on classical complement pathways, we examined synaptic termini at 10 dpi to determine if MDP treatment influences synapse elimination. Consistent with prior studies, infection with ZIKV leads to significantly reduced synapses (synaptophysin^+^homer1^+^ puncta) in the CA3 region of iMDP-treated mice, while administration of MDP beginning at 7 dpi completely prevented this reduction (Fig. 4A, B). Assessment of pre- and post-synaptic markers revealed significant reduction in homer1^+^, but not synaptophysin^+^, synaptic termini in only iMDP-treated ZIKV-infected mice at 10 dpi (Fig. 4B). Evaluation of synapses at 25 dpi revealed that administration of MDP during the acute phase of viral infection prevents not only the initial loss of synapses, but any additional synaptic elimination during viral clearance and recovery (Fig. 4C, D). Synaptic proteins within the CA1 region were also analysed to determine alterations due to ZIKV infection or MDP treatment and no differences were found (Fig. S4A, B). To assess if MDP treatment during acute infection impacted the generation of new neurons during recovery, animals were administered BrdU starting at 11 dpi to 14 dpi, and BrdU+ neurons in the DG of the HPC were quantitated at 25 dpi. At 25 dpi, ZIKV infection was associated with reduced numbers of newly mature neurons (NeuN+/BrdU+) and immature neurons (DCX+/NeuN+/BrdU+) within the SGZ of the HPC (Fig. 4 E, F), which was partially improved by MDP treatment (p value = 0.44, 0.43 respectively). These data indicate that MDP treatment critically impacts the extent of synapse elimination and limits reductions in newly born neurons during acute ZIKV infection of the CNS.

**Figure 4.**
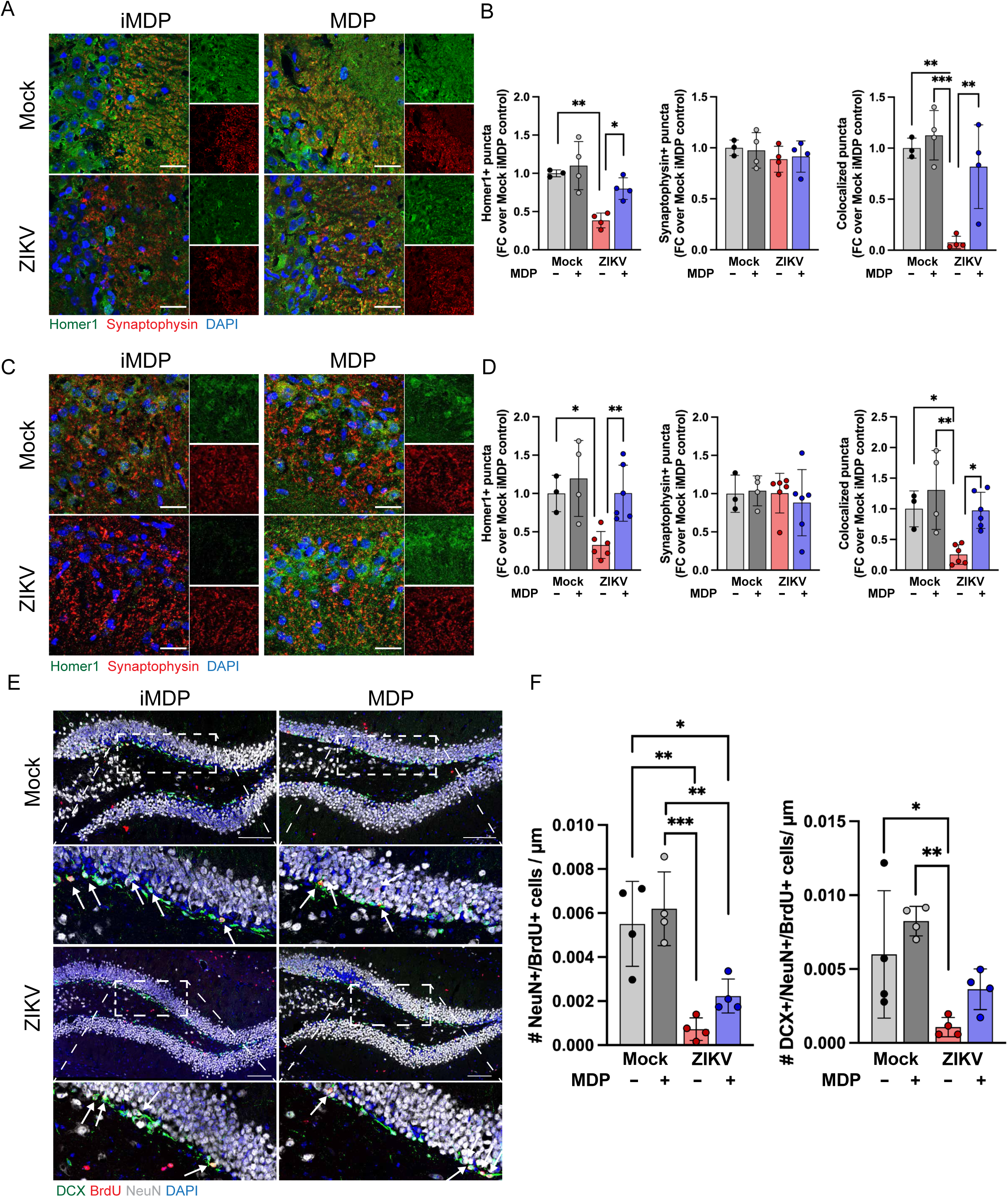
MDP attenuates loss of Homer1^+^ post-synaptic termini during recovery from ZIKV infection. (A) Representative images from ZIKV or mock-infected animals with either iMDP or MDP treatment, harvested at 10 dpi, and stained for detection of pre-synaptic (Synaptophysin, red) or post-synaptic (Homer1, green) termini in the CA3 region of the hippocampus. (B) Quantitation of Synaptophysin+, Homer1+, or co-localized puncta at 10 dpi. Mock iMDP n=3, Mock MDP n=4, ZIKV iMDP n=4, ZIKV MDP n=4. (C) Representative images from ZIKV or mock-infected animals with either iMDP or MDP treatment, harvested at 25 dpi, and stained for detection of pre-synaptic (Synaptophysin, red) or post-synaptic (Homer1, green) termini in the CA3 region of the hippocampus. (D) Quantitation of Synaptophysin+, Homer1+, or co-localized puncta at 25 dpi. Mock iMDP n=3, Mock MDP n=4, ZIKV iMDP n=6, ZIKV MDP n=6. (E) Representative images from ZIKV or mock-infected animals with either iMDP or MDP treatment, harvested at 25 dpi, and stained for detection of DCX (green), NeuN (grey), or BrdU (red), Location of insets are indicated in the image. DCX+/NeuN+/BrdU+ cells are indicated by arrows. (F) Quantification of NeuN+/BrdU+, and DCX+/NeuN+/BrdU+ cells within the dentate gyrus of the hippocampus at 25dpi. Mock iMDP n=4, Mock MDP n=4, ZIKV iMDP n=4, ZIKV MDP n=4. Scale bar, 25 µm (A, C), 100 µm (E). Each dot represents an individual mouse. Results are displayed as mean ± s.d. and were analyzed by two-way ANOVA, and corrected for multiple comparisons. All panels, \**p*<0.05, \*\**p*<0.01, *\*\*\*p<* 0.001.

### MDP-mediated preservation of synapses and increased phagocytosis requires NOD2 expression only by peripheral immune cells

As NOD2 is found in many non-immune and immune cell types including epithelial cells, platelets, neurons, astrocytes, T cells, macrophages, and microglia^47–52^, we utilized ZIKV-infected, iMDP- or MDP-treated bone marrow chimeric (BMC) mice to determine whether MDP-mediated synapse preservation and improved phagocytic capability requires NOD2 activation in peripherally derived versus CNS resident cellular targets. For these studies, we generated BMC mice in which *Nod2* is absent from the radiation sensitive bone marrow compartment (*Nod2-/-* bone marrow → irradiated WT recipients) or from radiation resistant CNS *Nod2-/-* cells (WT bone marrow → *Nod2-/-* recipients). WT bone marrow transplanted into irradiated WT recipients and *Nod2-/-* bone marrow transplanted into irradiated *Nod2-/-* recipients served as controls for effects of irradiation and engraftment. To preserve blood-brain barrier integrity and limit engraftment of donor cells into the CNS, all animals were irradiated with protective lead head shielding^53^. 5 weeks post-reconstitution, animals were infected with ZIKV i.c. followed by treatment with iMDP or MDP at 7 dpi, as per previous experiments. Chimerism did not alter weight loss, encephalitic scores, or survival (Fig. S4C - E). Congenic leukocyte markers CD45.1 (WT) and CD45.2 (*Nod2*-/-) were used to differentiate between donor and recipient cells via flow cytometry, revealing >75% engraftment (Fig. S4F). Only animals which were reconstituted with WT bone marrow and treated with MDP showed significant reductions in the percentage of Ly6C^HI^ cells (of the total monocyte population) in either blood or HPC samples (Fig. S4G) IHC assessment of synaptophysin+homer1+ puncta at 10 dpi within the CA3 region of the hippocampus revealed significant increases only in MDP-treated animals which were reconstituted with WT bone marrow (WT → WT recipient, WT → *Nod2-/-* recipient) (Fig 5A, B). To determine whether NOD2 activation impacts synapse engulfment by infiltrating macrophages or microglia, we quantified Homer1 puncta within Iba1+/TMEM119+ microglia versus Iba1+/TMEM119- macrophages within Lamp1+ lysosomes via Z-stack images from confocal microscopy with 3D mask rendering within the CA3 region using Imaris software(Fig. 5C). We measured the total volume of microglial and macrophage populations, including the Lamp1+ volume within each type of cell and the total number of Homer1+ puncta colocalized within macrophage or microglial lysosomes were also assessed(Fig. 5D, E). Total volume of microglial cells was assessed and we did observe a significant decrease of microglial volume in *Nod2-/-*recipient reconstituted with WT bone marrow after MDP treatment when compared to iMDP treated animals, yet not other pairing showed significant differences (Fig. 5E). MDP did, however, significantly increase the total volume of TMEM119-/Iba1+ cells in animals which received WT bone marrow (Fig. 5D). Similarly, Lamp1 volumes in microglia (within TMEM119+/Iba1+ masks) were not affected by chimerism or infection. However, Lamp1+ volume was significantly increased within TMEM119-/Iba1+ masks in MDP-treated samples, which received WT bone marrow, suggesting that MDP, acting through Nod2, increases the lysosomal activity of peripherally derived myeloid cells within the CNS during ZIKV infection. To determine if post-synaptic termini were being eliminated, we calculated the total number of Homer1+ puncta which were partially or fully engulfed within Lamp1+ masks either within microglia or macrophages. The total number of puncta were then normalized to the total volume of Lamp1 within each cell type to determine the rate at which each cell type was eliminating Homer1+ puncta. In both WT and Nod2-/- animals that received WT bone marrow and were treated with MDP, there was a significant reduction in the rate at which Homer1 was eliminated by microglial cells (TMEM119+/Iba1+; Fig. 5E). TMEM119-/Iba1+ cells showed no differences in Homer1 elimination across all groups suggesting that macrophages are not the main source of synapse elimination during ZIKV infection. Taken together, these data indicate that rescue of ZIKV-induced elimination of synapses, and increased phagocytic capacity induced by MDP is mediated via peripherally derived, infiltrating cells, with no contributions of CNS resident cells.

**Figure 5.**
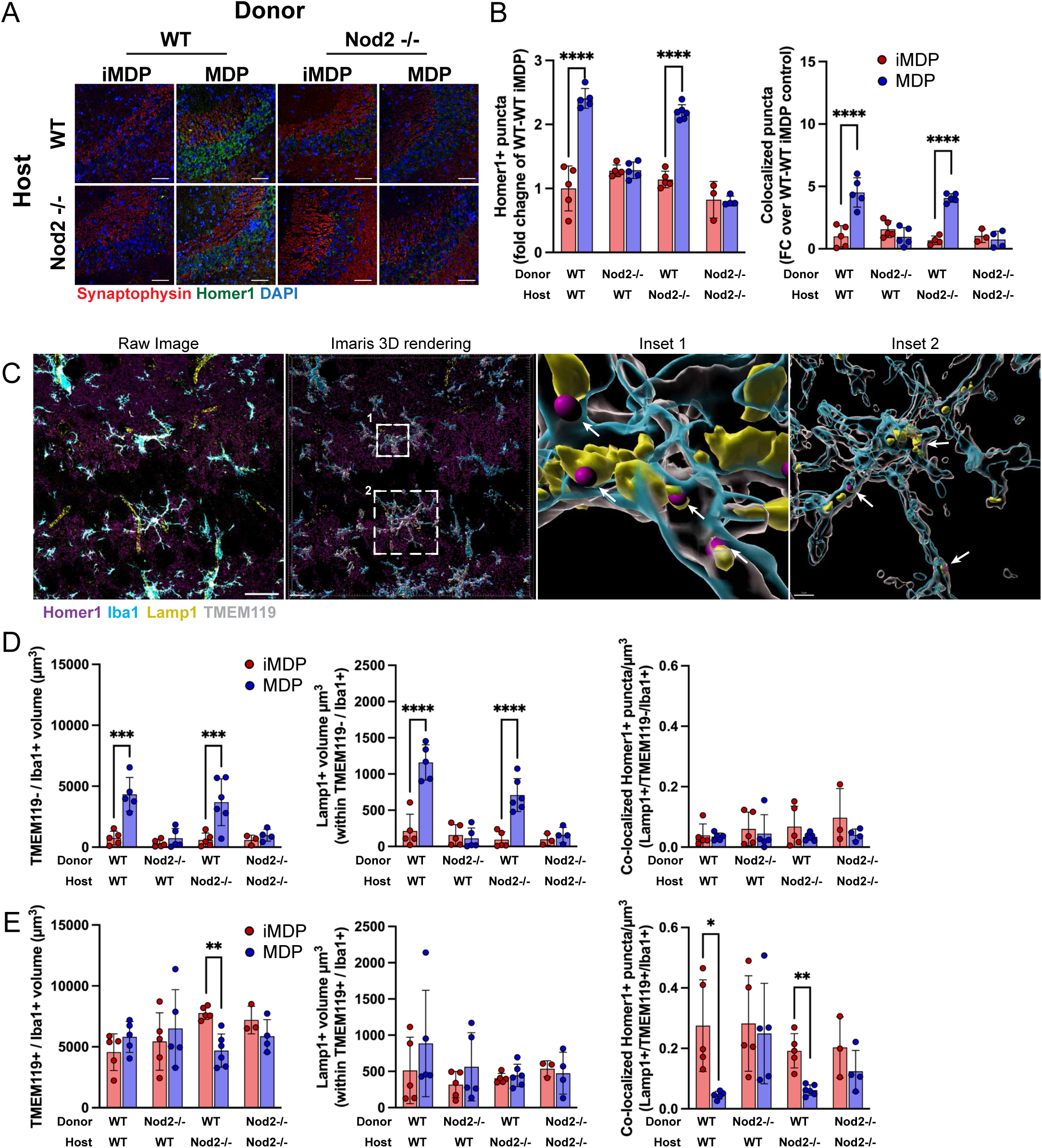
MDP-mediated alleviation of synapse loss during ZIKV encephalitis is dependent upon signaling through peripherally derived immune cells. (A) Representative immunostaining of hippocampal CA3 and (B) quantification of colocalized presynaptic and postsynaptic puncta using synaptophysin (red) and Homer-1 (green), respectively, at 10 dpi in generated bone marrow chimeric mice (WT donor – WT recipient (iMDP *n*=5, MDP *n*=5), *Nod2* KO donor – WT recipient (iMDP *n*=5, MDP *n*=5), WT donor – *Nod2* KO recipient (iMDP *n*=5, MDP *n*=6), *Nod2* KO donor - *Nod2* KO recipient (iMDP *n*=3, MDP *n*=4). (C) Representative imaging of hippocampal CA3 region for Homer1 (magenta), Iba1 (cyan), Lamp1 (yellow), and TMEM119 (grey). Imaris 3D rendering of the same image is shown with magnified insets labeled. Arrows indicate colocalized Homer1+ puncta within Lamp1+/Iba1+/TMEM119+ masks. (D, E) Quantitative analysis of macrophage (TMEM119-/Iba1+; D) and microglial (TMEM119+/Iba1+; E) volume, Lamp1+ volume within each cell type, and colocalized Homer1+ puncta with Lamp1+ masks within each cell type at 10 dpi. Data were pooled from two independent experiments. Scale bars, 25 µm (A, C; unless otherwise indicated in Imaris rendering). Data are displayed as mean ± s.d. and were analyzed by students’s t-test (B, D, E). All panels, \*\**p*<0.01, *\*\*\*p<* 0.001, \*\*\*\**p*< 0.0001.

## Discussion

Infiltration of inflammatory monocytes into the CNS during orthoflavivirus-induced encephalitis is necessary for effective virologic control and clearance^28,54^. However, their contribution to mechanisms that drive long-term cognitive consequences after viral resolution has not been examined. Using a murine model of ZIKV encephalitis that recapitulates learning and memory impairments that persist after viral clearance^34^, we found that short-term administration of the NOD2 agonist, MDP, during acute infection (7 – 10 dpi) converts infiltrating Ly6C^HI^ monocytes into a Ly6C^Lo^ phenotype with no impact on virologic control. Transcriptomic analyses of forebrain macrophages at 10 dpi revealed significantly increased M2 macrophage markers and phagocytic machinery, and decreases in inflammatory cytokine production. Using bone marrow chimera approaches, NOD2 activation via MDP was observed to selectively reprogram peripherally derived myeloid cells into a pro-resolution phenotype, enhancing their phagocytic capacity, alleviating hippocampal neuroinflammation, and preventing synapse elimination and loss of adult neurogenesis at 25 dpi, without requiring the involvement of resident CNS cells.

In mice, peripherally derived monocytes are generally divided into two principal subsets; Ly6C^Hi^ (CCR2^Hi^/CX3CR1^Lo^) inflammatory (M1) and Ly6C^Lo^ (CCR2^Lo^/CX3CR1^Hi^) anti-inflammatory (M2). Transcriptional profiling of immune cells from the forebrain after MDP treatment during the acute phase of ZIKV encephalitis revealed significant upregulation of M2-associated markers (*Arg1, Maf, Cpt1a, Grn, Tgfb1, Cx3cr1*) in macrophages with downregulation of M1 inflammatory associated genes (*Il1b, Ccr5, Ptgs2, Tlr2, Socs3, Cd86*) compared to iMDP-treated samples. However, we observed no differences in *Il10, Mrcl* (CD206), *Retnla, Chi3l3, Anxa1*, *Nr4a1*, *Klf2, Klf4, Pparg,* or*Tnfaip3* (A20) in the macrophage cluster suggesting that MDP is priming a shift to a more regulatory function without a complete alteration of phenotypic machinery. Our results are similar to what has been reported previously concerning MDP treatment and a subsequent polarization toward an M2b profile characterized by reduced inflammatory cytokines, elevated anti-inflammatory cytokine production, and increased phagocytic capacity^14,55^.

During infection with orthoflaviviruses, synapse elimination within the hippocampus is driven by activation of the classical complement cascade in which C1q localizes to synapses and promotes C3 deposition, thus tagging synaptic elements for elimination by microglia^32,56,57^. Cognitive impairment following recovery from flavivirus encephalitis is strongly linked to microglial-mediated synapse elimination with genetic ablation of complement (C3, C3aR) or CD8^+^ T cell-derived IFNγ signaling preserving synapse integrity and ameliorating memory deficits^32,34^. In contrast, infiltrating MDMs primarily support antiviral mechanisms, produce inflammatory cytokines and chemokines, and clear infected cells and debris, with less specificity for directly targeting synapses for elimination^28,58^. Consistent with this, MDM populations exhibited lack of Homer1+ puncta, even with increased phagocytic machinery due to MDP treatment. In contrast, MDP-treated mice exhibited significant reductions in colocalized Homer1+ puncta within microglia-associated lysosomes. However, bone marrow chimera studies showed that the effect of MDP requires NOD2 expression only in peripherally derived MDM, as only animals reconstituted with active NOD2 bone marrow and treated with MDP showed reductions in colocalized Homer1+ puncta within microglia-associated lysosomes.

Importantly, the persistence of post-synaptic termini within the CA3 region of the hippocampus in animals treated with MDP during ZIKV infection suggests that either these synapses are not targeted for elimination or that neurons within this region are actively producing signals that deter aberrant synaptic elimination. In our model, the latter explanation is more probable as we saw no alterations in *C1qa*, *C1qb, C3ar1* or *C3* expression, which is required for synapse elimination during CNS infection with orthoflaviviruses^32,45,56,59^, in the context of MDP treatment in any of our experiments. Instead, we observed decreased *Cd47* and *Tgfb* expression within the hippocampus in iMDP-treated animals, which is consistent with published findings demonstrating TGFβ, and it’s associated signaling, can be functionally linked to CD47 pathways and promote resistance to phagocytosis via transcriptional machinery that is conserved in neurons^60,61^. Finally, as TGFβ is required to maintain homeostatic, non-phagocytic microglial states^62^, limited activation of microglia would also reduce synapse elimination^34^. Thus, the combination of an overall reduced inflammatory milieu and upregulation of regulatory and anti-inflammatory molecules by MDMs during MDP treatment might allow neurons in the ZIKV-infected brain to retain CD47 to inhibit synapse engulfment.

Activation of NOD2 via MDP canonically induces the NFkB and MAPK signaling pathways, which have been shown to increase inflammatory cytokine expression^6,7^. However, recent studies indicate that the effects of MDP exposure on macrophage responsiveness and function depend on context and timing. Macrophage pre-exposure to MDP, or sustained NOD2 signaling, suppresses production of TNF, IL-6, and IL-12 and TLR-induced inflammation^14^. This is consistent with our transcriptomic findings showing reduced inflammatory cytokine expression and inflammatory signaling in macrophage populations after MDP administration. In our study, we also detected increased CD68, Lamp1, and phagocytic receptors, including *Mertk*, *Axl*, *Trem2*, in macrophage, but not microglial, populations after short-term treatment with MDP. However, post-synaptic proteins within the hippocampus were not eliminated, implicating an enhanced ability to clear pathologic debris without targeting synaptic structures. Similary, long-term MDP administration in a mouse model of Alzheimer’s disease similarly correlated with upregulation of molecules associated with phagocytosis (TREM2, CD68, LAMP2) in myeloid cells, which were linked to reduced amyloid burden and improved cognitive outcomes^55^. Our findings indicate that short-term administration of MDP increases phagocytic capacity of macrophages, yet reduces the overall inflammatory mileui within the ZIKV-infected hippocampus, preventing synapse elimination and loss of adult neurogenesis long-term.

T cell entry into the CNS during viral encephalitis, induced by chemoattractants including CXCL10, CXCL9, CCL5, CCL4, and CCL3, is necessary for viral resolution and the elimination of infected cells^63–68^. After MDP treatment, we found significant reductions in macrophage-specific expression of *Ccl3, Ccl4, Cxcl9,* and *Cxcl10* and a significant increase in *Ccl2* within the hippocampus. Single-cell RNAseq analysis did not identify any cell type with a corresponding significant increase in *Ccl2* expression, suggesting its upregulation within the hippocampus occurs within cell types not fully represented in the data set, including astrocytes and neurons, which can produce high levels of *Ccl2* under inflammatory conditions^69,70^. The increase in *Ccl2* with reductions in *Ccl3, Ccl4, Cxcl9,* and *Cxcl10* supports the observed shift from an acute inflammatory state, with reduced leukocyte recruitment, subsequent reductions in IFNγ levels, with selective maintenance or recruitment of M2-like monocytes/macrophages that assist in clearing cellular debris and facilitating repair^71–74^. This is subsequently supported by our data showing significantly reduced levels of *Ifng* mRNA and fewer CD8^+^ and CD4^+^ T cells within the hippocampus, as well as increased phagocytic capacity of macrophages due to MDP treatment.

In our study, we administered MDP at a time-point when viral titers peak within the brain (7 dpi)^34^ to model human presentation for clinical care. As Ly6C^Hi^ inflammatory monocytes/macrophages infiltrate the brain, they contribute to viral control and initiate adaptive mechanisms for viral clearance^54,75^. However, while administration of MDP acutely did not suppress antiviral immunity, it limited mechanisms known to induce long-term neurocognitive sequelae. MDP is a well-defined microbial ligand with known pharmacology and history as an immune adjuvant^76^, and therefore targeting NOD2 in monocyte/macrophage populations offers an attractive approach to reduce neuroinflammation-driven pathology. In the context of ZIKV-induced encephalitis, MDP-based immunomodulation may be used to mitigate innate immune-mediated CNS damage and improve neurologic outcomes without deleterious effects on virologic control.

## Limitations of the study

This study uncovers a role of peripherally derived macrophages in the induction of mechanisms that lead to long-term reductions of neural correlates of learning and memory. However, cell specific contributions to prolonged inflammation within the CNS during recovery from ZIKV encephalitis remains incompletely defined. Moreover, although we demonstrate that Nod2 signaling in peripherally derived cells is critical for beneficial effects of MDP treatment, we are still unable to determine the exact cell types responsible. Future work incorporating transgenic mice with cell-type specific ablation of Nod2 signaling will be essential to validate these findings. Additionally, this work would benefit from behavioral tests of spatial learning which would link MDP-mediated reductions in inflammatory mediators and attenuation of synapse elimination with functional outcomes relevant to human populations recovering from orthoflavivus induced encephalitis.

## Acknowledgments

The authors wish to acknowledge the Genome Technology Access Center, Washingon University School of Medicine, St. Louis, MO for their services in sequence alignment and quality control. Funding for this work was provided by a Canada Excellence Research Chair in Neurovirology and Neuroimmunology, Canada Foundation for Innovation John R. Evans Leaders Fund Award, US National Institutes of Health grant R01AI160188 (all to R.S.K), and an Ontario Graduate Scholarship (to A.H.W.D).

## Author contributions

J.D.H. and R.S.K conceived the project and wrote the manuscript, with input from all authors. J.D.H. performed all experiments and analyzed the data. A.D. analyzed the transcriptomics data. J.L. and M.D.B provided experimental assistance and contributed to data analysis. T.S.A provided essential scientific expertise. R.S.K was responsible for supervision and funding acquisition.

## Materials and Methods

### Animals

Male and female mice (8 weeks old) were used for each experiment. C57BL/6J (strain #000664), C57BL/6J CD45.1 (C57BL/6J-*Ptprc^em6Lutzy^*/J ; strain #033076), Nod2 KO (B6.129S1-*Nod2^tm1Flv^*/J; strain #005763) and Ccr2 KO (B6.129S4-Ccr2^tm1lfc^/J; strain #004999) were obtained from Jackson Laboratories. All experiments followed the guidelines approved by the Washington University School of Medicine Animal Safety Committee and Animal Care and Veterinary Services at Western University.

### ZIKV encephalitis model

The ZIKV-Dakar strain was used for intracranial infections. Mice were deeply anesthetized and intracranially administered 1 x 10^4^ focus-forming units (f.f.u.) of ZIKV-Dakar. Viruses were diluted in 10 µl of 0.5% fetal bovine serum (FBS) in Hank’s balanced salt solution (HBSS) and injected into the third ventricle of the brain with a guided 29-guage needle. Mock-infected mice were intracranially injected with 10 µl of 0.5% FBS in HBSS into the third ventricle of the brain with a guided 29-guage needle.

### Muramyldipeptide (MDP) administration

Mice were injected with muramyldipeptide (MDP; InvivoGen) diluted in saline (10 mg/kg) or the L-L inactive isomer of MDP diluted in saline (10 mg/kg), intraperitoneal (i.p.) daily starting on day 7 until day 10 post-infection.

### Viral burden – Focus forming assay

Vero cells were seeded in flat-bottom 96-well plates at 3 × 10^4^ cells/well in a volume of 100 μL per well. The next day, virus stocks or pre-weighed homogenized and clarified tissue supernatants were serially diluted in infection media (DMEM with 2% heat-inactivated FBS, 100 U/ml penicillin-streptomycin, and 10 mM HEPES). 100 μL of the diluted samples were added to Vero cell monolayers and incubated for 1 h at 37°C in 5% CO_2_. Subsequently, cells were overlaid with 100 μL of 1% methylcellulose in Minimum Essential Medium (MEM; Sigma #M0275) supplemented with 100 U/ml penicillin-streptomycin, 10 mM HEPES, and GlutaMAX (Gibco #35050–061). Plates were fixed 48h after virus inoculation. The methylcellulose overlay was first gently removed with a multichannel pipette and then, 300 μL of 4% PFA/PBS was added to each well for 1 h at room temperature. After three washes in PBS with 0.05% Tween 20 (Sigma #P1379) (PBS-T), samples were incubated on a plate rocker with 1 μg/mL of mouse anti-ZIKV monoclonal antibody (mAb) C60 diluted in permeabilization buffer (PBS, 0.1% saponin [Sigma #S7900], and 0.1% bovine serum albumin [BSA; Sigma #A2153]) for 2 h at room temperature or overnight at 4°C. Primary mAb was removed after three washes with PBS-T, and samples were incubated with secondary peroxidase-conjugated goat anti-mouse IgG (Sigma #A5278) diluted 1:500 in permeabilization buffer for 1 h at room temperature on a rocker. After three washes with PBS-T, virus-infected foci were developed using KPL TrueBlue peroxidase substrate (SeraCare #5510–0050), washed twice with Milli-Q water, and counted using an CTL-S6 Universal Analyzer (ImmunoSpot). Viral titer was expressed on a log10 scale as f.f.u. per gram of tissue.

### Immunohistochemistry and confocal microscopy

On the indicated day post-infection, mice were anesthetized and perfused with ice-cold PBS, followed by 4% PFA-PPS. Brain tissues were harvested and placed in 4% PFA-PBS for 24 hours. Tissues were washed in 3X with PBS, and then placed in 30% sucrose. Tissues were embedded by flash-freezing in OCT using dry ice. Tissues were sliced into 10 uM thick coronal sections using a cryostat at -20°C and mounted on SuperFrost Plus Slides. Coronal brain sections were washed with PBS, permeabilized with 0.3% Tween-20/Triton X-100 in PBS, and nonspecific antibody binding was blocked with 5% goat/donkey serum for 1 hour at room temperature (RT). Slides were washed 1x with PBS and incubated with the designated primary antibody for that experiment at 4°C overnight. After block, slides were exposed to primary antibodies overnight at 4 °C. The next day, after washing 3X with PBS, slides were incubated with appropriate secondary antibody for 1 hour at RT. Slides were washed 3X with PBS, nuclei were stained with DAPI (1:1000) for 5 min at RT followed by transferring the slides to sit in PBS for 5 minutes. Slides were coverslipped using ProLong Gold Antifade Mountant. Images were acquired on a Zeiss LSM 880 confocal laser scanning microscope and processed with accompanying software. Images on the Zeiss LSM 880 confocal laser scanning microscope were taken at 40 or 63X. Exposure settings for imaging during microscopy remained consistent for every image taken in that experiment.

### Antibodies for immunohistochemistry

The following primary antibodies were used for IHC analyses: IBA1(1:200, Invitrogen, PA518039), TMEM119(1:400, Cell Signaling, 90840S), CD68(1:200, BioRad, MCA1957), Lamp1(1:50, BD Pharminogen, 553792), Homer1(1:200, Synaptic Systems, 160 019), Synaptophysin(1:250, Synaptic Systems, 101 004), DCX(1:250, Cell Signaling, 4604S), BrdU(1:200, Abcam, ab6326), NeuN(1:500, Sigma Aldrich, ABN90P). Secondary antibodies conjugated to Alexa-405 (1:200, Invitrogen, cat # A-4258), Alexa-488 (Invitrogen, cat # A-21206, # A-78942), Alexa-555 (Invitrogen, cat # A-21435, # A-31572), or Alexa-647 (Invitrogen, cat # A-21447, # A-21450) were used at a 1:400 dilution.

### BrdU labeling

To perform in vivo BrdU labeling, mice were injected intraperitoneally every 12 h for 4 days with 50 mg per kg (body weight) BrdU (Sigma-Aldrich, B5002). To visualize BrdU accumulation in tissue slices, tissue was prepared as described above in Immunofluorescence microscopy.

Tissue sections were incubated for 5 min in distilled water. DNA was denatured by incubating sections in ice-cold 1 N HCl for 10 min at 4 °C, followed by incubation in 2 N HCl at 37 °C for 30 min. Acid was neutralized by washing sections in 0.1 M borate buffer twice, followed by three washes in PBS. Slides were blocked with 5% donkey serum and 0.1% Triton X-100 in PBS for 1 h at 20–21 °C. After blocking, slides were washed once with PBS and incubated with rat anti-BrdU (1:200) overnight at 4 °C. Staining was performed as described above.

### Image analysis

All image acquisitions and analyses were performed blinded. For image quantification, a minimum of two to three sections per animal were averaged to obtain one biological replicate. For synaptic terminals, FIJI was used to threshold single-plane confocal images taken at 63 X magnification and the number of homer1+ or synaptophysin+ puncta were quantified.

Overlapping pixels positive for both homer1 and synaptophysin within the CA3 region were considered colocalized and quantified separately. A minimum of 9 images were counted per animal (3 images per section, minimum 3 sections per mouse). For percent area measurements, blinded images were thresholded to the same value for each channel in FIJI and measured. For cell number quantification (DCX, BrdU, and NeuN), in FIJI, images were set to the same contrast settings, and the number of positive cells was manually counted by a blinded individual. Cell number was divided by the length of the SGZ of the DG in µm for each image. Three to four sections were quantified per animal. For 3D image analysis, Imaris software was used (Version 11.0, Bitplane AG, Zurich, Switzerland). Confocal z-stacks were taken at 63 x magnification and individual channels were thresholded to the same value. Masks were generated for TMEM119, Iba1, and Lamp1 positive stains. Iba1 positive masks were separated via colocalization with TMEM119 to generate two separate groups; Iba1+/TMEM119+ (microglia) and Iba1+/TMEM119- (macrophages). Lamp1 masks which colocalized within <0.1µm of microglial or macrophage masks were considered cell specific Lamp1. Homer1 puncta were determined via the spots function in Imaris and individual puncta were counted. Colocalization of homer1 puncta was determined by puncta that were <0.1µm from Lamp1 masks and total numbers of puncta colocalized with Lamp1 per cell type were assessed. A minimum of two images per section and two sections per animal were counted.

### Flow cytometry

Cells from the hippocampus and blood of mice were isolated at indicated d.p.i. and stained with fluorescence-conjugated antibodies (described below). Briefly, mice were anesthetized and perfused with ice-cold Dulbecco’s PBS (Gibco). Blood (100µl) was collected directly from the heart prior to perfusion, collected in tubes pre-coated with EDTA, and immediately placed on ice. ACK lysis buffer was then added to each blood sample for 10 minutes at RT while protected from light. An equal volume of PBS was added to halt the ACK buffer and samples were spun down at 2000 x *g* for 5 min. Excess ACK/PBS was aspirated from samples and the pellet was resuspended in 1ml PBS. Hippocampal brain tissue was dissected out, minced, and enzymatically digested at room temperature for 1 h with shaking. The digestion buffer contained collagenase D (Sigma, 50 mg ml^−1^), TLCK trypsin inhibitor (Sigma, 100 μg ml^−1^), DNase I (Sigma, 100 U μl^−1^), HEPES buffer, pH 7.2 (Gibco, 1 M) in HBSS (Gibco). The tissue was then pushed through a 70-μm strainer and pelleted by a 500 × *g* spin cycle for 10 min. To remove myelin debris, cells were resuspended in 37% Percoll and spun at 1,200 × g for 30 min. Cells were then resuspended in FACS buffer. TruStain fcX anti-mouse CD16/32 (BioLegend, cat. no. 101320, clone 93) was used to block the cells for 5 min at 4 °C, followed by cell surface staining for 15 min at 4 °C. Cells were washed twice in FACS buffer, fixed with 4% PFA, and resuspended in 2% PFA for data acquisition. Data were collected using a Fortessa X-20 instrument and analyzed with the software Flowjo.

### Antibodies for flow cytometry

The following antibodies were used for flow cytometry: CD45(BUV737, BD Biosciences, 748371), CD11b(BV421, Biolegend, 101235 ), CD4(PerCP/Cy5.5, Biolegend, 100433), CD8(BV711, Biolegend, 100747 ), CD103(BV605, Biolegend, 121433), CD69(PE/Cy7, Biolegend, 104512), Ly6C(AF700, Biolegend, 128024), Ly6G(PE/Cy7, Biolegend, 127617), Ly6G(BV605, Biolegend, 127639), CD115(BV711, Biolegend, 135515), NK1.1(FITC, Biolegend, 108717), CD45.1(BUV737, CytekBio, 367-0453-80), CD45.2(PE/Cy7, CytekBio, 60-0454-U100), P2RY12(PE, Biolegend, 848004), CD19(FITC, ThermoFisher, 11-0193-82).

### scRNA-seq

Mice (2 groups, *n*=2 per group, 3 mice pooled per *n*) were deeply anesthetized and perfused intracardially with ice-cold dPBS (Gibco). Forebrain tissue was aseptically dissected, minced, and enzymatically digested in HBSS (Gibco) containing collagenase D (Sigma, 50mg/ml), TLCK trypsin inhibitor (Sigma, 100μg/ml), DNase I (Sigma, 100U/μl), and HEPES 7.2 (Gibco, 1M), for 1h at 37°C while shaking. The tissue was pushed through a 70-μm strainer and spun down at 500*g* for 10min. To remove myelin debris, cells were resuspended in 37% Percoll and spun at 1200g for 30 min. To minimize cell loss and aggregation, cells were resuspended at 100 cells/μL in PBS containing 0.04% (w/v) BSA. ∼17,500 cells were partitioned into nanoliter-scale Gel Bead-In-EMulsions (GEMs) to achieve single-cell resolution for a maximum of 10,000 individual cells/sample. Poly-adenylated mRNA from an individual cell was tagged with a unique 16 basepair 10x barcode and 10 basepair unique molecular identifier utilizing the v2 Chromium Single Cell 3′ Library Kit and Chromium instrument (10x Genomics). Full-length cDNA was amplified to generate sufficient mass for library construction. cDNA amplicon size (∼400 basepair) for the library was optimized using enzymatic fragmentation and size selection. The final library was sequence-ready and contained four unique sample indexes. The concentration of the 10x single cell library was determined via qPCR (Kapa Biosystems). The libraries were normalized, pooled, and sequenced on the HiSeq4000 platform (Illumina). Four single-cell libraries were sequenced across an entire HiSeq4000 flow cell targeting ∼45,000 reads per cell.

All bioinformatics analysis and associated data processing, unless otherwise specified, was performed using the R statistical computing software (v4.2.2)^77^ and tidyverse (v2.0.0)^78^.

Seurat (v4.3)^79^ was used for analysis of the scRNAseq dataset unless otherwise specified. To exclude barcodes with abnormal counts or number of features, the number of counts and the number of features were log10-transformed and fitted to a B-spline curve using the ‘bs’ function with default parameters. Only barcodes that fall within the 99% prediction interval and those with less than 10% mitochondrial transcripts were included for downstream analysis. Samples in each experimental group were merged, then normalized on a per-cell basis using SCTransform (v0.4.2)^80^ with the top 3000 genes selected as variable features. Then, samples in the two experimental groups were integrated using reciprocal principal component analysis (RPCA) with default parameters implemented in Seurat. Principal component analysis (PCA) was then performed on the integrated dataset, followed by t-distributed stochastic neighbor embedding (t-SNE) calculation using the first 24 principal components (PCs) for visualization. To identify distinct cell types within the integrated dataset, using Seurat, a shared nearest-neighbor graph was generated using the first 24 PCs with default parameters, and clusters were assigned with the Louvain algorithm at a resolution of 0.3. By comparing the gene expression in each cluster to all other clusters using the Wilcoxon rank sum test with default parameters, distinct markers for each cluster were identified and used to annotate each cluster with reference to Tabula Muris^81^. Subclustering of selected major cell types were then performed using the Louvain algorithm at a resolution of 0.1.

### DE analysis and pathway enrichment

To correct for potential biases due to imbalanced barcode (cell) counts in each sample, a sample with high counts was downsampled to the average number of cells in other samples randomly without replacement. The Wilcoxon rank sum test with default parameters was used to identify differentially expressed genes (DEGs) in each cluster across experimental conditions.

Bonferroni correction was applied to the p-values to account for multiple comparisons. DEGs were defined as genes with log2-transformed fold-change less than -0.3 or over 0.3 and adjusted p-value less than 0.05. Over-representation analysis using gproflier2^82^ was performed for each cluster using the corresponding upregulated and downregulated DEGs separately to identify upregulated and downregulated pathways. Pathways from Gene Ontology^83,84^ biological processes and Reactome^85^ were used as references. After adjustment of p-values by Bonferroni correction, significant pathways with term sizes between 15 and 1000 were visualized as networks using EnrichmentMap (v3.5.0)^86^ in Cytoscape (v3.10.3)^87^ and clustered using the Markov Cluster Algorithm (MCL) implemented by AutoAnnotate (v1.5.2)^88^ using default parameters. Cluster annotations were manually assigned referencing Gene Ontology and Reactome databases.

### Cell-cell communication inference

LIANA+ (v1.7.1)^89^ and Python (v3.11)^90^ was used to determine putative interactions among immune cells in the scRNA-seq dataset. Gene symbols in the ‘consensus’ resource^91^ included in LIANA+ was first converted to mouse gene symbols using human-mouse ortholog data provided by The HUGO Gene Nomenclature Committee (HGNC) Comparison of Orthology Predictions (HCOP). To establish a baseline of cell-cell communications between forebrain immune cells in the context of ZIKV encephalitis, only cells isolated from ZIKV-infected mice that received the iMDP treatment were included in the cell-cell communication inference. The counts from these cells were scaled such that each cell has 1000 total counts, then log-normalization was performed on the scaled counts. Using the ‘consensus’ resource, cell-cell communications were calculated using CellPhoneDB^92^, Connectome^93^, log2FC, NATMI^94^, and SingleCellSingalR^95^, then the predictions from all methods were ranked and aggregated. Only ligand-receptor predictions with specificity rank < 0.05 were considered for downstream analysis. A ligand-receptor prediction was considered altered if either the ligand or the receptor was a differentially expressed gene. ‘Log2FC’ of a ligand-receptor interaction was defined as the average of the log2-transformed fold change of the ligand and receptor in the respective cell types.

### Real-time quantitative RT–PCR

Total RNA was extracted using *RNeasy*® Plus Mini kit (Qiagen) as per manufacturer’s instructions. Total RNA was converted to cDNA using high-capacity cDNA Reverse Transcription kit (Thermo Fisher Scientific). qRT-PCR was carried out using *Power* SYBR PCR Master Mix (Applied Biosystems, cat. no. 4367659) with target-specific primers. Specific primers for mouse immune targets (listed in table 5) were purchased from Sigma Aldrich and analyses were performed using the CFX384 Touch RT-PCR detection system (Bio Rad). Data for murine samples were normalized to GAPDH. qPCR analysis is represented as fold change using the 2^-ΔΔCt^ method as compared to Day 0 animals. Statistics for these studies were performed on ΔC_T_ values.

### Generation of Bone Marrow Chimeras

Chimeric mice were generated using slight modifications to a previously published protocol^96^. Bone marrow ablation was achieved by irradiating four- to five-week-old wild type C57BL/J (CD45.1) and Nod2 KO (CD45.2) mice with 10-10.5 Gy (X-ray) irradiation while protecting the skull via shield (Precision X-Ray Irradiation). The next day, bone marrow cells were collected from tibia, femur, and humerus of five-week old C57BL/6J (CD45.1) or Nod2 KO (CD45.2) animals. Each irradiated, recipient mouse was administered 5 x 10^6^ sex-matched bone marrow cells via retro-orbital injection. Five weeks after bone marrow transplantation, mice were infected with ZIKV-Dakar or mock solution via intra-cranial injection as outlined above.

Chimerism of each animal was confirmed by flow cytometric analysis of peripheral blood cells as well as immune cells harvested from brain tissue. Single cell staining procedures used were exactly as outlined above in “Flow Cytometry”. Cells were analyzed using a BD FACSCanto II (BD Biosciences) cytometer. All analyses were conducted using FlowJo software (v11, BD Life Sciences).

### Statistical analyses

For all analyses, data was processed randomly, and quantification was performed by experimenters who were blinded to experimental groups. All values are expressed as mean± SD. Statistical significance was assigned when *p* values were <0.05 using Prism version 10 (GraphPad). Tests, number of animals (n), mean values, and statistical comparison groups are indicated in the Figure legends. Statistics for qPCR data were performed on ΔC_T_ values.

## Supplemental Information

Document S1. Figures S1-S5

Table S1. scRNAseq markers per cell type for identification

Table S2. scRNAseq DEGs per cell type

Table S3. Number of significant DEGs per cell type

Table S4. scRNAseq enriched pathways per cell type

Table S5. qPCR primer sequences

---

**Figure S1:**
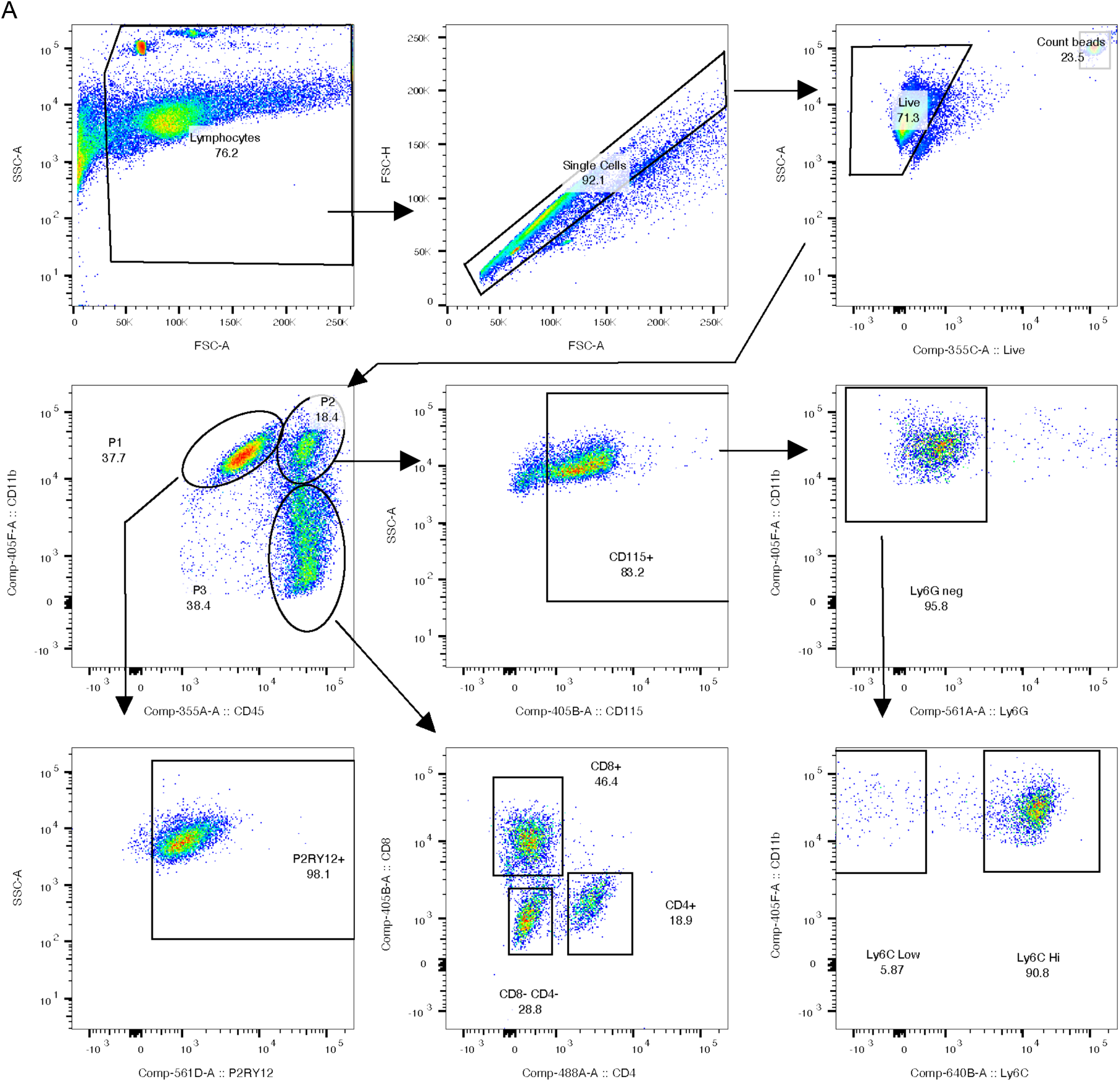
Representative flow cytometry gating strategy (Linked to Figure 1). Representative plots illustrating the hierarchical gating strategy used to isolate monocyte, microglial, and T cell populations. Initial events were gated on a Forward Scatter (FSC-A) vs. Side Scatter (SSC-A) plot to exclude debris and isolate the leukocyte population. Doublets were excluded by comparing Forward Scatter Area (FSC-A) to Forward Scatter Height (FSC-H). Viable cells were identified by the exclusion of a Fixable Viability Dye (FVD). From the live, single-cell population, cells were gated on CD11b+ and CD45+ parameters, generating three distinct subsets; CD45^MID^/CD11b^HI^ (P1), CD45^HI^/CD11b^HI^ (P2), and CD45^HI^/CD11b- (P3). P1 cells were further gated to assess microglial cells via P2RY12+ staining. Monocytes/macrophages were characterized by selecting cells positive for CD115, then negative gating against Ly6G to remove neutrophils, and then assessing the levels of Ly6C. P3 cells were gated against CD8 and CD4 to determine cellular subsets. Fluorescence minus one (FMO) controls were used to set gates. Numbers within or adjacent to gated regions represent the percentage of the parent population.

**Figure S2:**
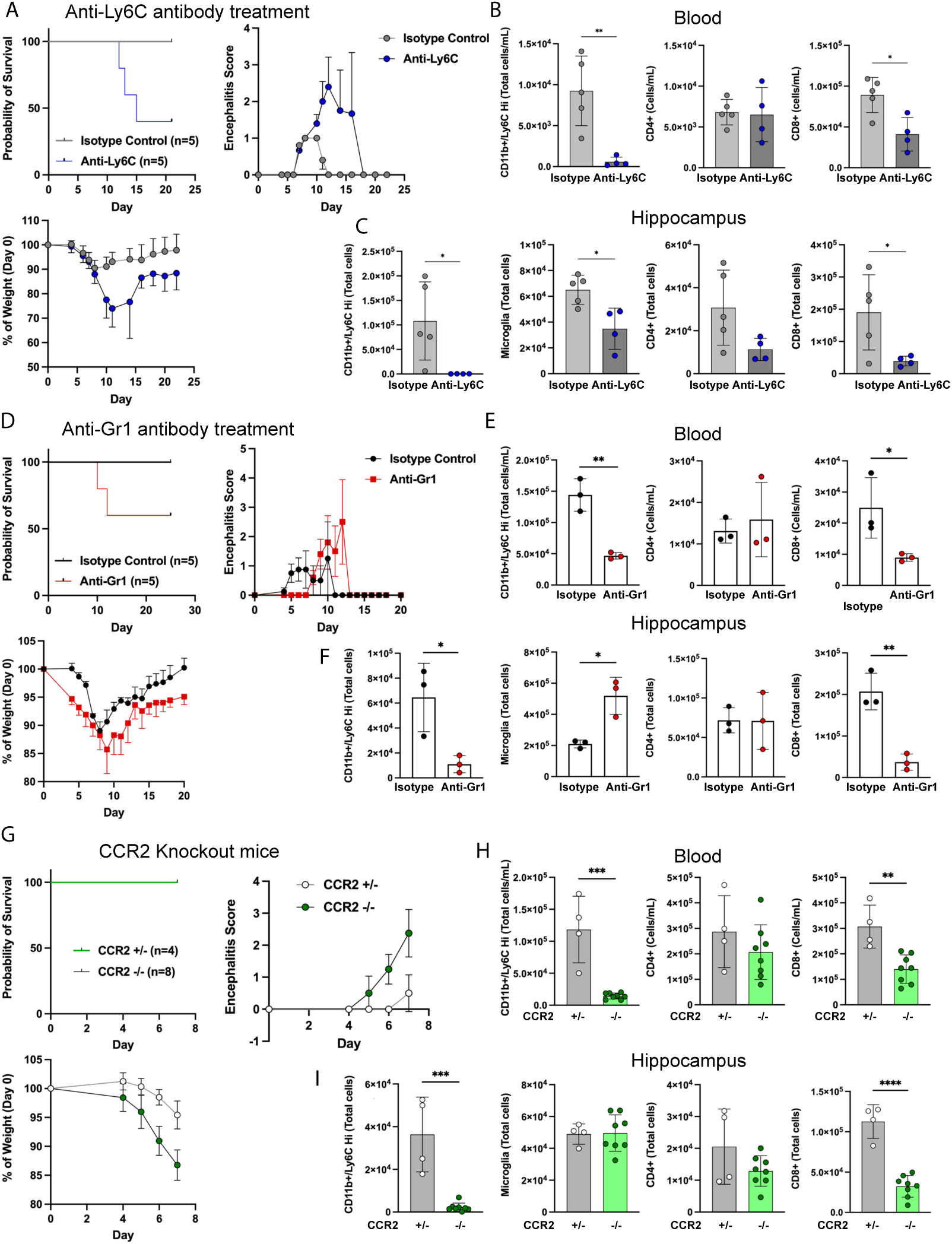
Alternative methods utilized to reduce infiltration of peripheral monocytes (Linked to Figure 1). (A-I) Numerous alternative methods to deplete monocyte involvement during intracranial ZIKV infection were assessed. (A) 8 week-old C57BL/6J animals were inoculated intracranially with ZIKV-Dakar (1x10^4^ FFU) and administered anti-Ly6C (Monts1) or isotype control (Rat IgG2a; 100µg/injection, retro-orbital injections given 48hours apart starting at time of infection [Day 0]). Animals were harvested at 25 dpi (Isotype, n=5; Anti-Ly6C n=5). Survival, encephalitic score, and % of weight change were assessed. Samples from (B) blood and (C) hippocampi were taken from a separate cohort of animals (Isotype, n=5; Anti-Ly6C n=5) at 7 dpi to determine differences in monocytes, T cell subsets (CD4^+^, CD8^+^), and microglia (hippocampus only). (D) 8 week-old C57BL/6J animals were inoculated intracranially with ZIKV-Dakar (1x10^4^ FFU) and administered anti-Gr1 or isotype control (Rat IgG2b; 200µg/injection, retro-orbital injections given 72hours apart starting on 3 dpi, 3 injections total). Animals were harvested at 25 dpi (Isotype, n=5; Anti-Ly6C n=5). Survival, encephalitic score, and % of weight change were assessed. Samples from (E) blood and (F) hippocampi were taken from a separate cohort of animals (Isotype, n=3; Anti-Ly6C n=3) at 7 dpi to determine differences in monocytes, T cell subsets (CD4^+^, CD8^+^), and microglia (hippocampus only). (G) 8 week-old Ccr2+/- or Ccr2-/- animals were inoculated intracranially with ZIKV-Dakar (1x10^4^ FFU) Animals were harvested at 7 dpi (Ccr2+/-, n=4; Ccr2-/- n=8). Survival, encephalitic score, and % of weight change were assessed. Samples from (H) blood and (I) hippocampi were taken to determine differences in monocytes, T cell subsets (CD4^+^, CD8^+^), and microglia (hippocampus only). Each dot represents an individual mouse. Data are displayed as mean ± s.d. and were analyzed by students’s t-test. All panels, \**p*<0.05, \*\**p*<0.01, *\*\*\*p<* 0.001, \*\*\*\**p*< 0.0001.

**Figure S3:**
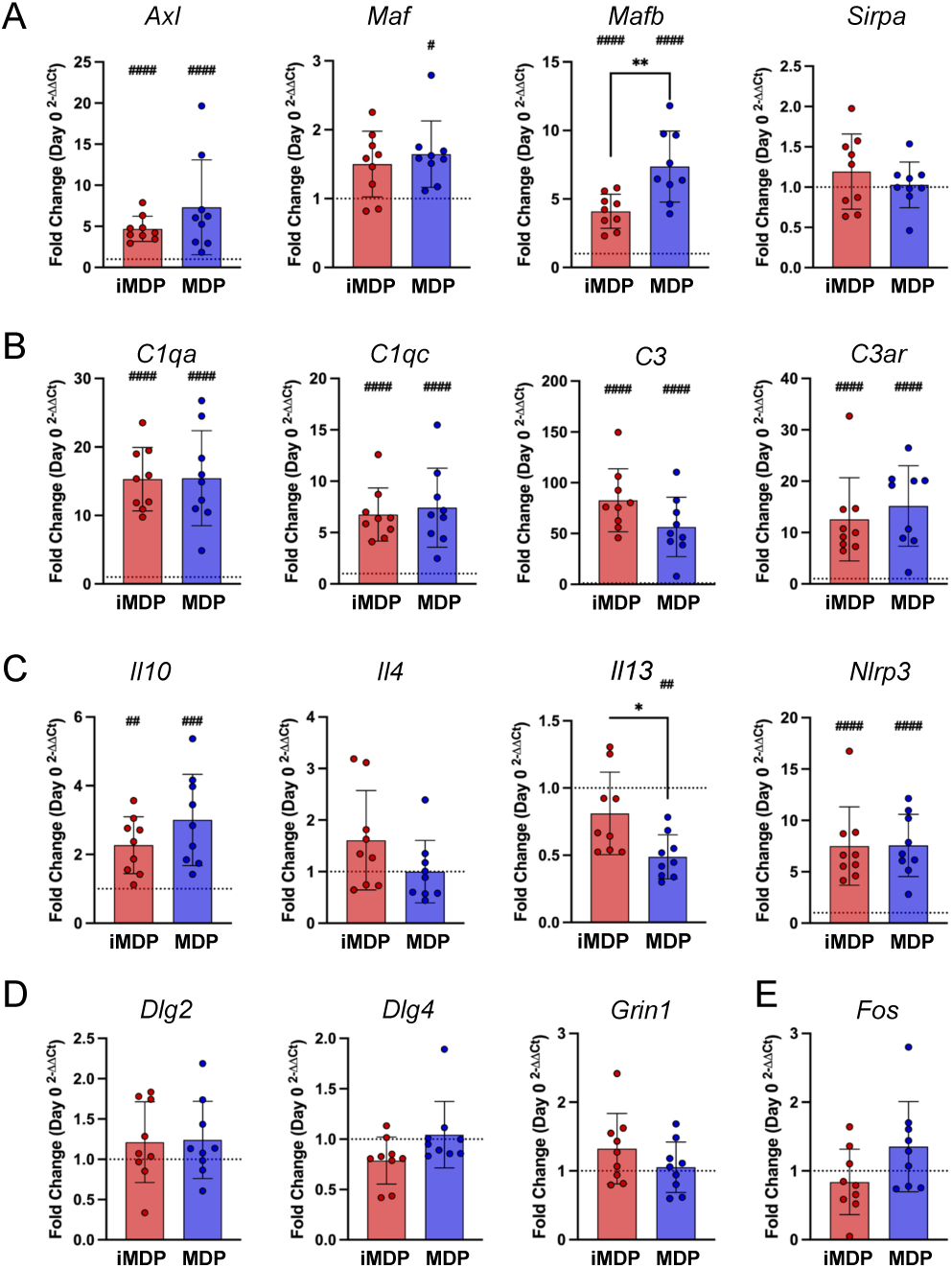
Extended qPCR data from hippocampi of iMDP or MDP-treated mice (Linked to Figure 2). (A-E) qPCR analysis of (A) complement activation, (B) cytokines and inflammasome signaling, (C) genes associated with phagocytosis, (D) synaptic maintenance, and (E) activity dependent transcription. Each dot represents an individual mouse (A-E, Day 0 n=6, iMDP n=9, MDP n=9). Data are presented as fold change of Day 0 samples (mean ± s.d.). Dotted lines on graphs represent fold change of Day 0 samples (1). Asterisks denote statistical significance between iMDP and MDP treatments (\**p*<0.05, \*\**p*<0.01, \*\*\**p*<0.001). Hash signs denote statistical significance between Day 0 and treatment (#*p*<0.05, ##*p*<0.01, ###*p*<0.001, ####*p*<0.0001). Statistical analysis was performed using one-way ANOVA with multiple comparisons (A-E).

**Figure S4:**
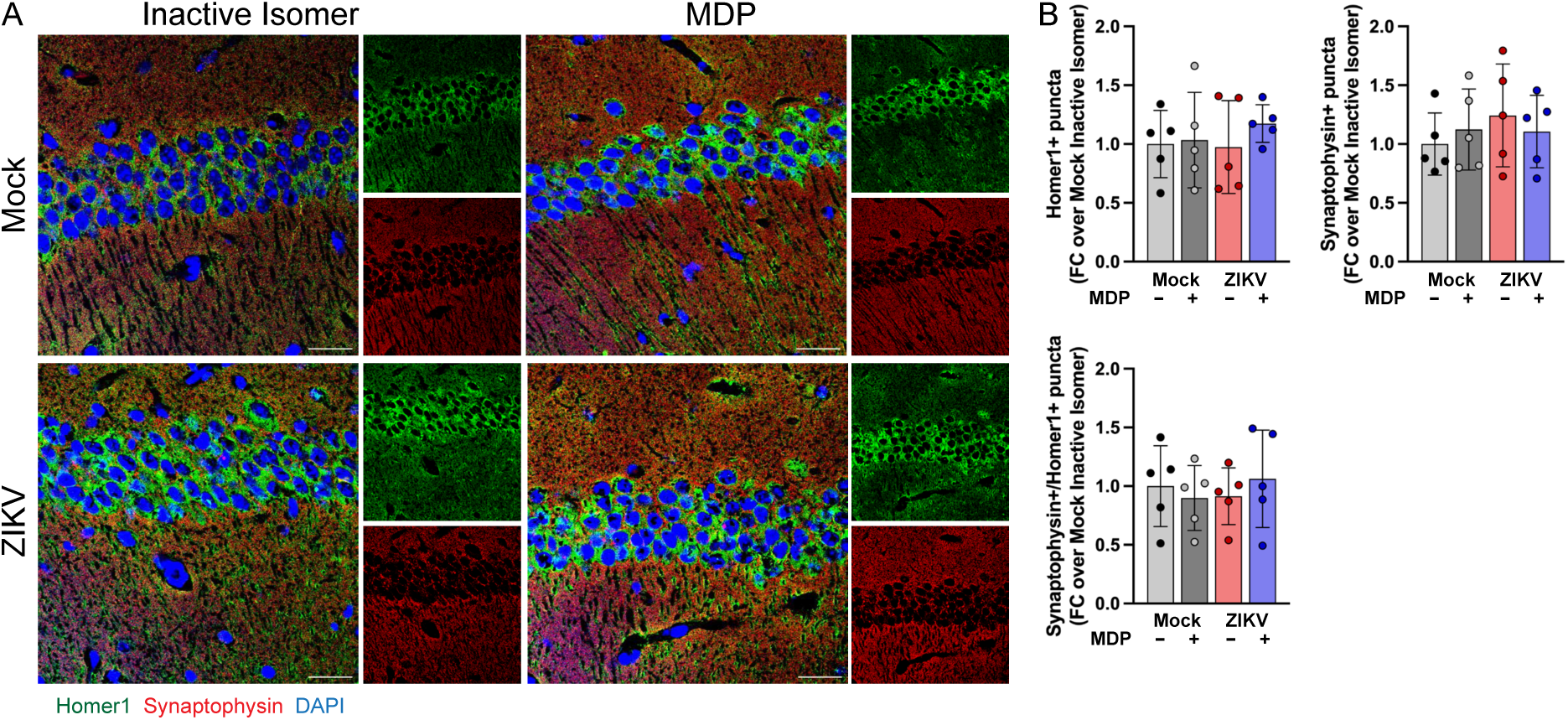
ZIKV encephalitis does not alter synaptic protein levels or synapses in the CA1 region of the hippocampus (Linked to Figure 4). (A-B) 8-week-old C57BL/6J animals were inoculated intracranially with ZIKV-Dakar (1x10^4^ FFU) and administered an inactive isomer of MDP (iMDP, 10mg/kg) or MDP (10mg/kg) i.p. starting at day 7 post-infection and given daily injections until harvest (10 dpi). (A) Representative images from ZIKV or mock-infected animals with either iMDP or MDP treatment, harvested at 10 dpi, and stained for detection of pre-synaptic (Synaptophysin, red) or post-synaptic (Homer1, green) termini in the CA3 region of the hippocampus. (B) Quantitation of Synaptophysin+, Homer1+, or co-localized puncta at 10 dpi. Scale bar, 25 µm. Each dot represents an individual mouse; n=5 per group. Results are displayed as mean ± s.d. of fold change of mock, iMDP-treated animals. Raw puncta values were used for statistical analysis and analyzed by two-way ANOVA, corrected for multiple comparisons.

**Figure S5:**
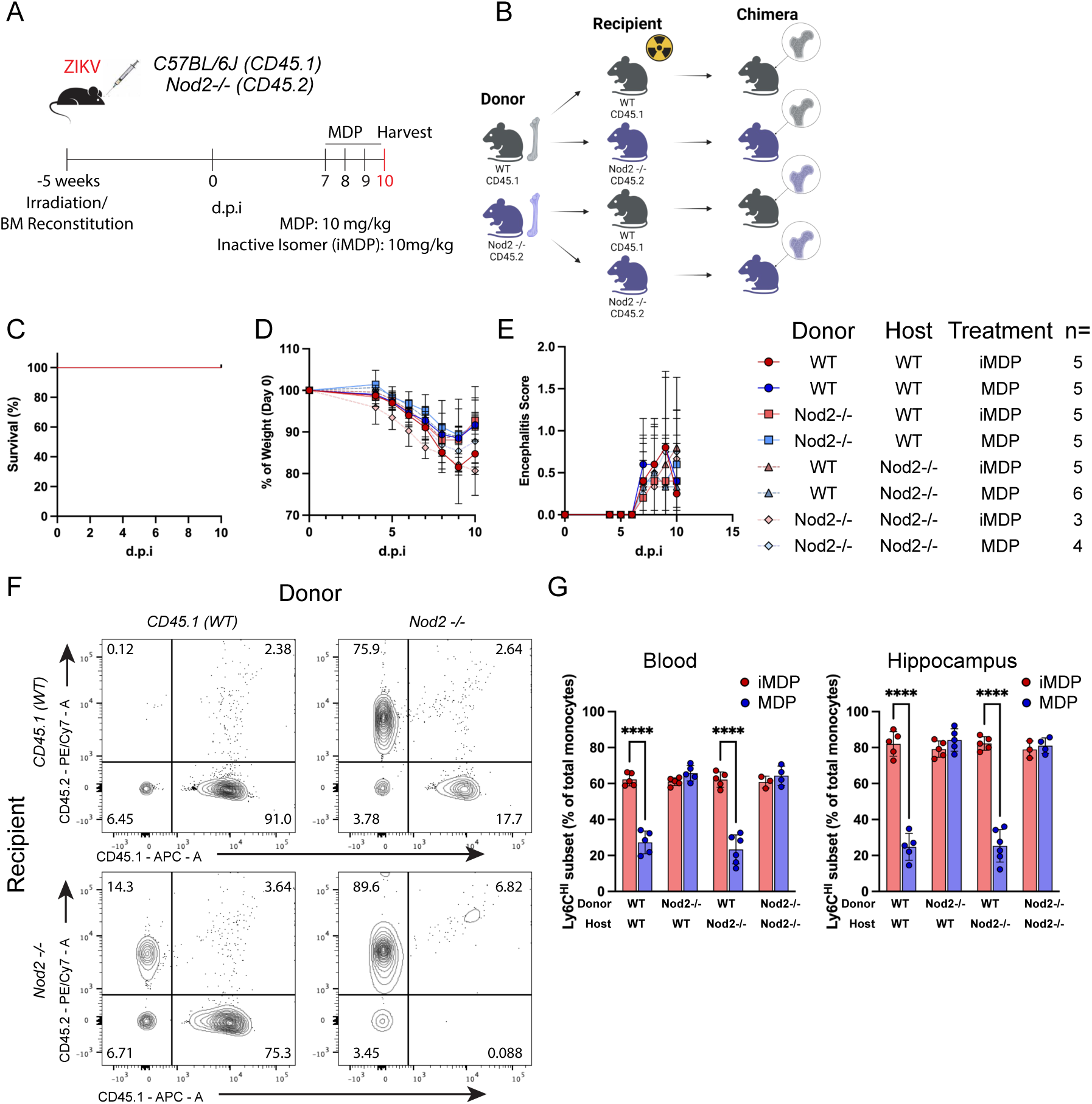
BM chimera generation and assessment (Linked to Figure 5). (A-B) (A) Experimental scheme for generating wild-type and *Nod2* KO bone marrow chimeric mice. Five weeks after irradiation and reconstitution with donor bone marrow hematopoietic stem cells, recipient mice were inoculated intra-cranially with 1x10^4^ FFU of ZIKV-Dakar. Mice were then treated with MDP or inactive isomer control (iMDP) beginning at 7 dpi and tissue was harvested at 10 dpi. (B) Graphical illustration of chimerism generated (WT donor – WT recipient (iMDP *n*=5, MDP *n*=5), *Nod2* KO donor – WT recipient (iMDP *n*=5, MDP *n*=5), WT donor – *Nod2* KO recipient (iMDP *n*=5, MDP *n*=6), *Nod2* KO donor - *Nod2* KO recipient (iMDP *n*=3, MDP *n*=4). (C) Chimerism did not induce differences in lethality due to ZIKV encephalitis. (D) Relative weight change was not significantly altered due to chimerism or treatment. Data is expressed as the percentage of Day 0 body weight. (E) Encephalitc scores were not significantly different due to chimerism or treatment. (F) Representative flow cytometry plots are shown analyzing the percentage of CD45.1+ and CD45.2+ cells of peripheral blood from recipient WT-CD45.1+ cells reconstituted with donor *Nod2* KO CD45.2+ cells and recipient *Nod2* KO CD45.2+ cells reconstituted with donor WT-CD45.1+ cells. (G) Percentage of Ly6C^HI^ cells in the monocyte population within blood or hippocampi of chimeric mice at 10 dpi. Each dot represents an individual mouse. Data are pooled from two independent experiments. Data are displayed as mean ± s.d. and were analyzed by students’s t-test (G). All panels, \*\*\*\**p*< 0.0001.

